# Spatial and temporal PCP protein dynamics coordinate cell intercalation during neural tube closure

**DOI:** 10.1101/278499

**Authors:** Mitchell T. Butler, John B. Wallingford

## Abstract

Planar cell polarity (PCP) controls the convergent extension cell movements that drive axis elongation in all vertebrates. Though asymmetric localization of core PCP proteins is central to their function, we currently understand little about PCP protein localization as it relates to the subcellular behaviors that drive convergent extension. Here, we have used high magnification time-lapse imaging to simultaneously monitor cell intercalation behaviors and the localization of the PCP proteins Prickle2 and Vangl2. We observed the expected asymmetric enrichment of PCP proteins, but more interestingly, we also observed tight temporal and spatial correlation of PCP protein enrichment with contractile behavior in cell-cell junctions. These patterns of localization were associated with similar pattern of protein turnover at junctions as assessed by FRAP. In fact, dynamic enrichment of PCP proteins was linked more strongly to junction behavior than to spatial orientation. Finally, recruitment of Prickle2 and Vangl2 to cell-cell junctions was temporally and spatially coordinated with planar polarized oscillations of actomyosin enrichment, and all of these dynamic relationships were disrupted when PCP signaling was manipulated. Together, these results provide a dynamic and quantitative view of PCP protein localization during convergent extension and suggest a complex and intimate link between the dynamic localization of core PCP proteins, actomyosin assembly, and polarized junction shrinking during cell intercalation of the closing vertebrate neural tube.

## Introduction

Convergent extension (CE) is the evolutionarily-conserved morphogenetic engine that drives elongation of the body axis in animals ranging from insects to mammals (Tada and Heisenberg, 2012). Multiple cell behaviors can contribute to CE, but by far the most well understood process is cell intercalation, by which cells rearrange in a polarized manner (Walck-Shannon and Hardin, 2014). Cell intercalation, in turn, is thought to be driven by multiple subcellular behaviors, including extension of mediolaterally-directed cellular protrusions (e.g. (Shih and Keller, 1992) and active shrinkage of mediolaterally oriented cell-cell junctions (e.g. (Bertet et al., 2004; Blankenship et al., 2006). More recent data suggest that these two subcellular behaviors likely work in concert (Sun et al., 2017; Williams et al., 2014). Understanding the molecular mechanisms governing protrusive activity and junction shrinking during cell intercalation will be essential to understanding convergent extension.

The junction shrinking mechanism for cell intercalation was initially described in *Drosophila* (Bertet et al., 2004; Blankenship et al., 2006) and was subsequently identified in both epithelial and mesenchymal cells in vertebrates (Lienkamp et al., 2012; Nishimura et al., 2012; Shindo and Wallingford, 2014; Trichas et al., 2012). In all tissues examined by live imaging, junction shrinkage is accompanied by pulsed actomyosin contractions that are restricted to or enriched at mediolaterally-oriented cell-cell junctions and absent from or less common at the junctions perpendicular to the anterior-posterior axis (Bertet et al., 2004; Blankenship et al., 2006; Shindo and Wallingford, 2014; Williams et al., 2014). A major unresolved question concerns the molecular mechanism by which actomyosin activity is restricted to specific cell-cell junctions during intercalation.

In *Drosophila*, pair-rule genes and Toll receptors are crucial regulators of polarized actomyosin (Pare et al., 2014; Zallen and Wieschaus, 2004), but homologous genes do not appear to be involved in vertebrates. Instead, the most well characterized regulator of cell intercalation in vertebrates is the Planar Cell Polarity (PCP) signaling system (Butler and Wallingford, 2017). Indeed, PCP signaling controls cell intercalation during gastrulation and neural tube closure in frogs, fish, and mice, and mutations in PCP genes are now a well-defined genetic risk factor for human neural tube birth defects (NTDs)(De Marco et al., 2014; Juriloff and Harris, 2012; Wallingford et al., 2013). Understanding PCP signaling is therefore a critical challenge in developmental cell biology.

One fundamental principle of PCP signaling is that cell polarity is imparted by asymmetric enrichment of the core PCP proteins (Strutt and Strutt, 2009). As shown first in *Drosophila*, the Prickle (Pk) and Van Gogh (Vangl) proteins act in concert on one side of the cell, and Dishevelled and Frizzled act on the complementary side (Axelrod, 2001; Bastock et al., 2003; Strutt, 2001; Tree et al., 2002). Interestingly, FRAP studies in *Drosophila* have shown that these patterns of enrichment are driven by planar polarization of the junctional turnover kinetics of PCP proteins, underscoring the dynamic nature of the PCP signaling system (Strutt et al., 2011). Similar patterns of enrichment and dynamic turnover have been reported in vertebrate multiciliated cells (Butler and Wallingford, 2015; Chien et al., 2015; Shi et al., 2016), but less is known about PCP protein localization during cell intercalation.

For example, complementary, asymmetric domains of PCP protein enrichment have been described during vertebrate CE (Ciruna et al., 2006; Jiang et al., 2005; McGreevy et al., 2015; Ossipova et al., 2015; Roszko et al., 2015; Yin et al., 2008), but how PCP protein enrichment is coordinated in space and time with the subcellular behaviors that drive intercalation remains essentially unexplored. This gap in our knowledge is critical, because recent work demonstrates that PCP proteins are required for the junction shrinking behaviors that contribute critically to cell intercalation, especially in epithelial cells (Lienkamp et al., 2012; Nishimura et al., 2012; Shindo and Wallingford, 2014). Thus, there is a pressing need for a quantitative, dynamic picture of PCP protein localization as it relates both to subceullar behaviors involved in cell intercalation and to the actomyosin machinery that drives them.

To this end, we established methods for robust quantification of PCP protein localization in a living vertebrate neural plate as well as methods for correlating PCP protein dynamics with the subcellular behaviors that drive epithelial cell intercalation. Strikingly, we find that in addition to expected patterns of spatial asymmetry, PCP protein enrichment is tightly linked to cell-cell junction behavior: Prickle2 (Pk2) and Vangl2 were dynamically enriched specifically at shrinking cell-cell junctions and depleted from elongating junctions during cell intercalation. FRAP analysis revealed that these patterns of enrichment reflected differences in the kinetics of protein turnover at junctions. Moreover, Pk2 enrichment was temporally and spatially correlated with planar polarized oscillations of junctional actomyosin enrichment. Importantly, all of these dynamic relationships are disrupted when PCP signaling is manipulated. Thus, our studies reveal an intimate link between the dynamic localization of core PCP proteins, actomyosin assembly, and polarized junction shrinking during cell intercalation of the closing vertebrate neural tube.

## Results

We characterized PCP protein dynamics in the neural plate of *Xenopus*, as studies in this animal consistently prefigure similar results in mammalian systems yet provide exceptional views of dynamic cell biological processes (Kieserman et al., 2010). Because Dishevelled and Frizzled function in both PCP and canonical Wnt signaling, their roles are more difficult to interrogate, so we focused instead on Prickle and Vangl, which act solely in PCP signaling and are required for neural tube closure in vertebrates, including *Xenopus* (Darken et al., 2002; Goto et al., 2005; Goto and Keller, 2002; Kibar et al., 2001; Takeuchi et al., 2003). Previous work suggests that Prickle and Vangl localize to the anterior face of cells in the *Xenopus* neural plate (Ossipova et al., 2015), but we sought to establish a more robust quantification of this pattern as a foundation for the time-lapse studies described below.

GFP-Vangl2 displayed a strong asymmetric bias to anterior cell faces (Fig. 1A), consistent with the previous report (Ossipova et al., 2015). Core PCP proteins are encoded by multi-gene families, and despite the reported role of Pk1 in convergent extension, GFP-Pk1 did not display asymmetric localization. On the other hand, Prickle2 (Butler and Wallingford, 2015) was strongly enriched anteriorly (Fig. 1A; Supp Fig. 1 related to Fig. 1). Knockdown experiments confirmed that Pk2 was physiologically relevant for convergent extension and neural tube closure in *Xenopus* (Supp. Fig. 1, related to Fig. 1). In double-labeling experiments, GFP-Vangl2 strongly co-localized with RFP-Pk2 (Fig. 1A).

**Figure 1:**
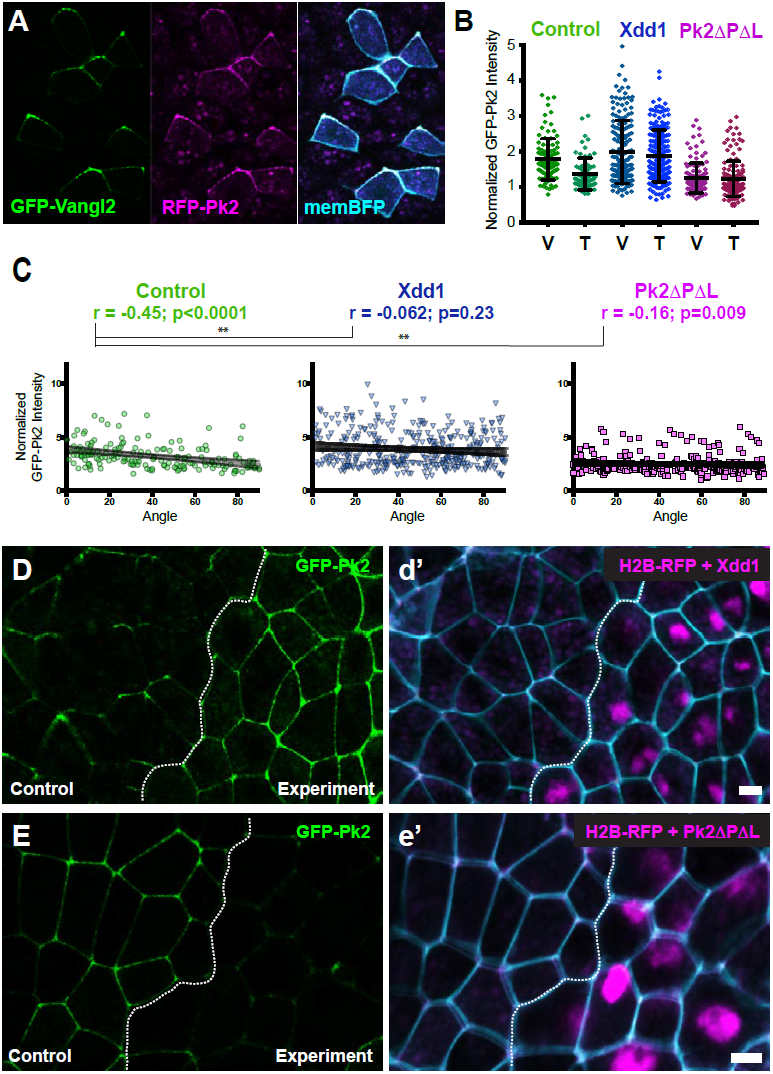
Planar polarized localization of Prickle2 and Vangl2. **(A)** Neural epithelium mosaically labeled with GFP-Vangl2, RFP-Pk2, and membraneBFP showing the overlapping localizations of Pk2 and Vangl2. Anterior is up. **(B)** Graph plotting GFP-Pk2 intensity along V junctions (0-45° relative to mediolateral axis) and T junctions (46-90° relative to mediolateral axis) normalized as a ratio to the cytoplasmic intensity in control cells (n=101 V, 71 T) and cells expressing Xdd1 (n=171 V, 199 T) and Pk2-ΔPΔL (n=128 V, 142 T). Ctrl V vs. T, p<0.0001; Xdd1 V vs. T, p=0.5799; Pk2-ΔPΔL V vs T, p=0.173; Ctrl V vs Xdd1 T, p=0.5770; Ctrl T vs Pk2-ΔPΔL V, p=0.0268. **(C)** Distributions of normalized GFP-Pk2 intensity plotted against the angle of the junction at which the intensity was measured in control cells and cells expressing Xdd1 or Pk2-ΔPΔL. Correlation coefficients for Xdd1 and Pk2-ΔPΔL were significantly different from controls using the Fischer R-to-Z transformation. n=172 junctions (Control), n=263 junctions (Xdd1) and n=245 junctions (Pk2-ΔPΔL) from 4 experiments, 5 embryos (Xdd1); 3 experiments, 7 embryos (Pk2-ΔPΔL). **(D-E)** Confocal images of *Xenopus* neural epithelia labeled evenly with Myl9-GFP and membraneBFP and mosaically with H2B-RFP, serving as a tracer for either Xdd1 (C) or Pk2-ΔPΔL (D). Scale = 10 μm.

We quantified these localization patterns at the population level by binning cell-cell junctions into two groups based on their orientation. To maintain a consistent nomenclature with previous work on cell intercalation in other systems, we refer to junctions aligned within 45 degrees of the mediolateral axis as “V-junctions” and the perpendicular junctions (between 45 and 90 degrees off the mediolateral axis) as “T-junctions.” An important subtlety to note here is that the V-*junctions* are *mediolaterally* aligned, but actually separate *cells* that are *anteroposterior* neighbors. We found that GFP-Pk2 was significantly enriched at V-junctions as compared to T-junctions (Fig. 1B). We also quantified these data at the level of individual junctions, and we observed a significant correlation between GFP-Pk2 pixel intensity and junction angle, with higher enrichment along more mediolaterally-oriented junctions (Fig. 1C, green).

To test our quantification schemes, we took advantage of the fact that in many other systems, disruption of any one core PCP protein leads to loss of polarized enrichment of the others. We used targeted injection to co-express reagents for disrupting PCP signaling together with a nuclear RFP lineage tracer, allowing us to compare normal and experimental cells in the same embryo (Fig. 1D, E). Using both ensemble and individual metrics, we detected significant disruption of Prickle2 asymmetries in cells expressing of the well defined, PCP-specific dominant negative of Dvl, Xdd1 (Sokol, 1996; Wallingford et al., 2000)(Fig. 1B, C blue, pink). Similar results were obtained when we expressed the dominant-negative form of Prickle2 lacking the PET and LIM domains (Pk2-ΔPΔL; (Butler and Wallingford, 2015; Takeuchi et al., 2003)(Fig. 1B, C, blue, pink). These results establish the *Xenopus* neural plate as an effective, quantitative platform for studies of PCP protein localization.

### Epithelial convergent extension in the closing *Xenopus* neural tube involves PCP-dependent polarized junction shrinking

Convergent extension is an inherently dynamic process, as cells constantly exchange neighbors. Tissues engaged in convergent extension therefore differ markedly from the settings in which PCP protein localization has most commonly been studied. We therefore sought to exploit the strengths of *Xenopus* embryos to image PCP protein dynamics together with the subcellular behaviors that drive convergent extension in the closing neural tube. Curiously, the neural plate of *Xenopus* consists of two cell layers, an outer epithelial layer and a deeper mesenchymal layer (Schroeder, 1970). While cell intercalation of the deep mesenchymal cells has been characterized (Elul et al., 1997; Keller et al., 1992), the behaviors of the overlying epithelial cells have not. This distinction is not trivial because it is the outer epithelial cells that display the robust patterns of PCP protein localization described above. To understand how PCP protein localization relates to convergent extension cell behaviors, we had first to characterize cell behaviors in this epithelium.

To this end, we used confocal microscopy and image tiling to generate high-magnification time-lapse movies of the neural epithelium of *Xenopus* embryos (see methods). Traces of individual cells in these movies (Fig. 2A, B) closely resemble those of previous studies (Keller et al., 1992), but the image tiling employed here provided sufficient magnification to also quantify the behavior of individual cells. Tracking of cell clusters revealed extensive cell intercalations that were mediolaterally biased, resulting in convergence and extension (Fig. 2C).

**Figure 2:**
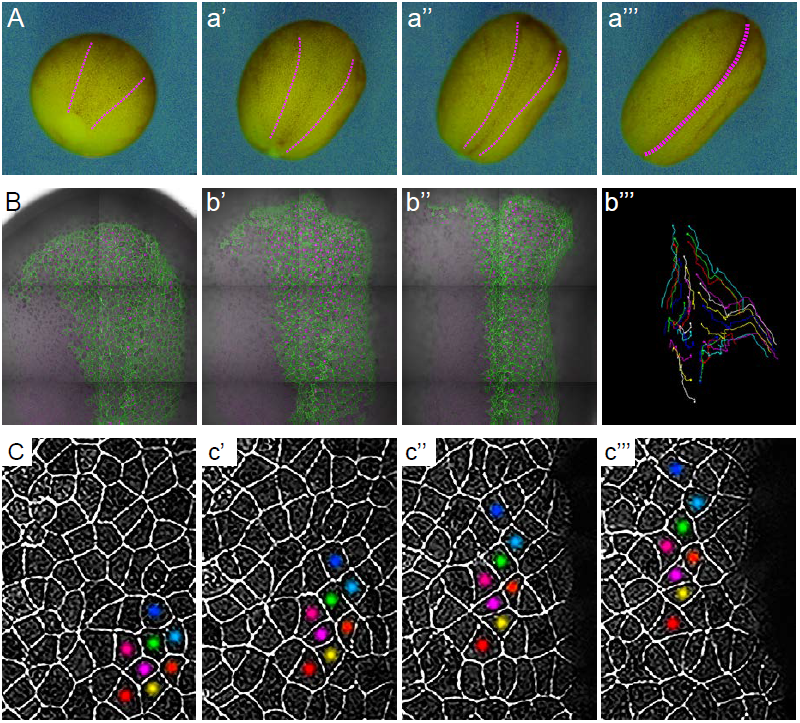
High magnification time-lapse imaging of convergent extension in the closing *Xenopus* neural tube. **(A)** Stereo image stills from a time-lapse movie of *Xenopus* neural tube morphogenesis from stages 12 to 19. **(B)** Stills from a time-lapse movie of the dorsal side of an embryo from stages 12 to 16. Confocal images of cells labeled with membraneGFP and H2B-RFP are overlain on the DIC image of the same embryo. B’’’ shows tracks cell movements over the duration of the movie. **(C)** Higher magnification images of cell rearrangements in neural ectoderm from the time-lapse shown in (B). Labeling of individual cells with colored dots across time points demonstrates the narrowing and lengthening of tissue.

Cell intercalations in the neural plate were associated with so-called T1 transitions, which are characterized by preferential contraction of junctions aligned in the mediolateral axis (Fig. 3A, a’), followed by elongation of new junctions perpendicularly along the anteroposterior axis (Fig. 3a’, a”)(Bertet et al., 2004). As above, we first quantified these behaviors at the ensemble level and found that mediolaterally oriented V-junctions preferentially shrank, while the perpendicular T-junctions elongated (Fig.3B). For a more granular view of these data, we also plotted the change in the length of cell-cell junctions against the average angle of that junction for all cells examined (n > 250). This analysis revealed a significantly positive correlation, with V-junctions preferentially shrinking along the mediolateral axis and T-junctions elongating perpendicularly (Fig. 3C). Finally, we also observed planar polarized formation and resolution of multicellular rosettes (Fig. 3D-F), as have been described in other epithelia (Blankenship et al., 2006; Lienkamp et al., 2012; Trichas et al., 2012).

**Figure 3:**
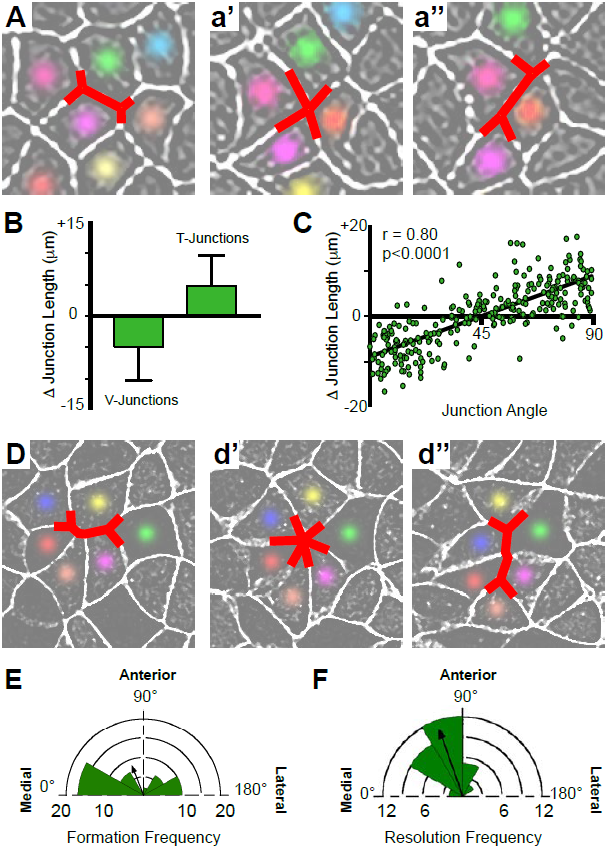
Polarized apical junction dynamics facilitate mediolateral cell intercalations. **(A)** Confocal images of junction dynamics in neural ectoderm labeled with membraneGFP. Red lines mark the shrinking of v-junctions during a T1 transition; after complete shrinkage mediolaterally **(a’)**, a new junction elongates perpendicularly along the AP axis **(a”)**. **(B)** Graph showing the mean change (+/- s.d.) in junction length for V- and T-junctions. **(C)** Plot of the average angles of junctions over 1800 seconds against the natural log of the fold increase or decrease in length over the same time frame. Each dot represents on cell-cell junction. n=267 junctions from 3 embryos across 3 different experiments. **(D)** The simultaneous mediolateral shrinking of several neighboring v-junctions (labeled in red) leads to formation of a multicellular rosette **(d’)**, and new junctions that emerge from the resolving rosette are oriented along the AP axis **(d’’)**. **(E)** Rose diagram plotting the orientation of shrinking junctions that lead to the formation of multicellular rosettes. n=42 junctions from 4 embryos across 3 separate experiments. **(F)** Rose diagram plotting the orientation of new junctions emerging from resolving rosettes. n=36 new junctions from 3 embryos across 3 separate experiments.

Because PCP signaling is essential for neural convergent extension (Wallingford and Harland, 2001; Wallingford and Harland, 2002), we next assessed the effect of PCP disruption specifically on junction shrinking behaviors in the neural epithelium. Using the mosaic approach described above, we found that expression of Xdd1 elicited the expected tissue-autonomous defect in the medially-directed movement of neural fold (Fig. 4A, magenta indicates Xdd1 expressing cells).

**Figure 4:**
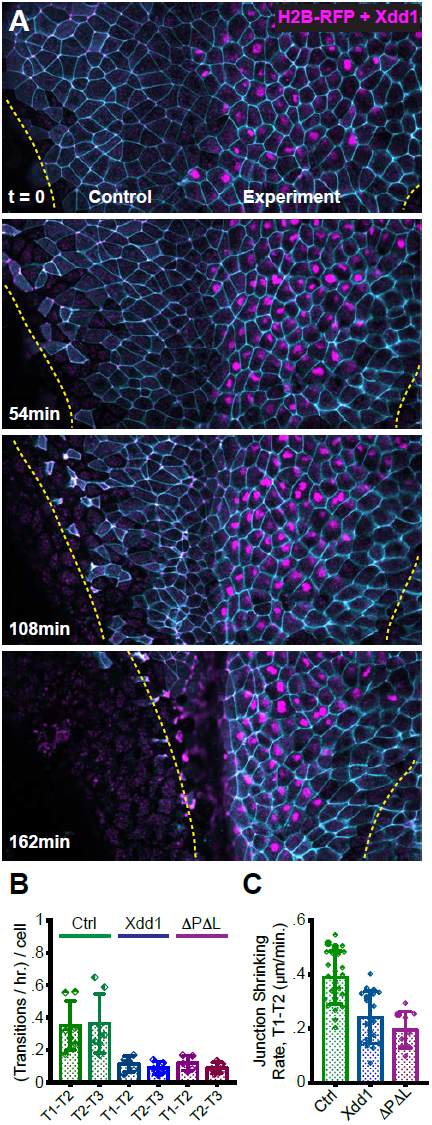
PCP function is required for polarized junction shrinking in the neural plate. **(A)** Confocal images from a *Xenopus* neural plate evenly labeled with membraneBFP and Utr-RFP, but mosaically co-expressing H2B-RFP (magenta) together with Xdd1 on one side (right). The lateral boundaries of the neural plate are marked in each frame by yellow lines, demonstrating that Xdd1 expression disrupts the medial movement of the right neural fold in comparison to the control fold on the left. **(B)** Graph with the total number of T1 transition events expressed as transitions per hour per cell. For statistical analysis, Control vs. Xdd1, and Control vs. Pk2-ΔPΔL ML, p=0.0025** for both classes of transitions (Mann-Whitney Test for significance). n=4 experiments, 7 embryos, 1,167 cells (Control); 3 experiments, 5 embryos, 560 cells (Xdd1); 3 experiments, 5 embryos, 841 cells (Pk2-ΔPΔL) **(C)** The calculated rate of junction contraction for completed Type 1 to Type 2 transitions (complete contraction of a v-junction, see Figure 2B). Ctrl vs. Xdd1, p<0.0001****, Ctrl vs. Pk2-ΔPΔL, p<0.0001****, Xdd1 vs. Pk2-ΔPΔL, p=0.2051ns. (Mann-Whitney statistical test). n=24 (Ctrl), n=19 (Xdd1), and n=9 (Pk2-ΔPΔL) junctions.

Analysis of individual cell behaviors in these embryos revealed that Xdd1 expression significantly reduced the number of productive T1 transitions in the neural plate (Fig. 3B, blue). Moreover, even the few productive T1 transitions that were observed in these embryos were significantly slowed (Fig. 4C, blue). Similar results were obtained with the dominant-negative Pk2-ΔPΔL (Fig. 4B, C, pink). Together with our data on PCP protein localization (above), these results establish the *Xenopus* neural plate as an effective platform with which to probe the relationship between epithelial cell intercalation behaviors and core PCP protein dynamics.

### Prickle2 and Vangl2 are dynamically enriched specifically at shrinking cell-cell junctions

With these imaging and analysis systems in place, we performed time-lapse imaging with an eye toward understanding the dynamic relationship between PCP protein localization and epithelial cell behaviors. This analysis revealed several novel insights. First, we found that GFP-Pk2 was dynamically enriched at shrinking junctions and likewise depleted from elongating junctions over time (Fig. 5A). Moreover, instantaneous changes in GFP-Pk2 intensity were strongly correlated to simultaneous changes in junction length (Fig. 5B, b’). Strikingly, the correlation between GFP-Pk2 intensity and junction behavior was substantially stronger than that reported in Fig. 1 for junction orientation.

**Figure 5:**
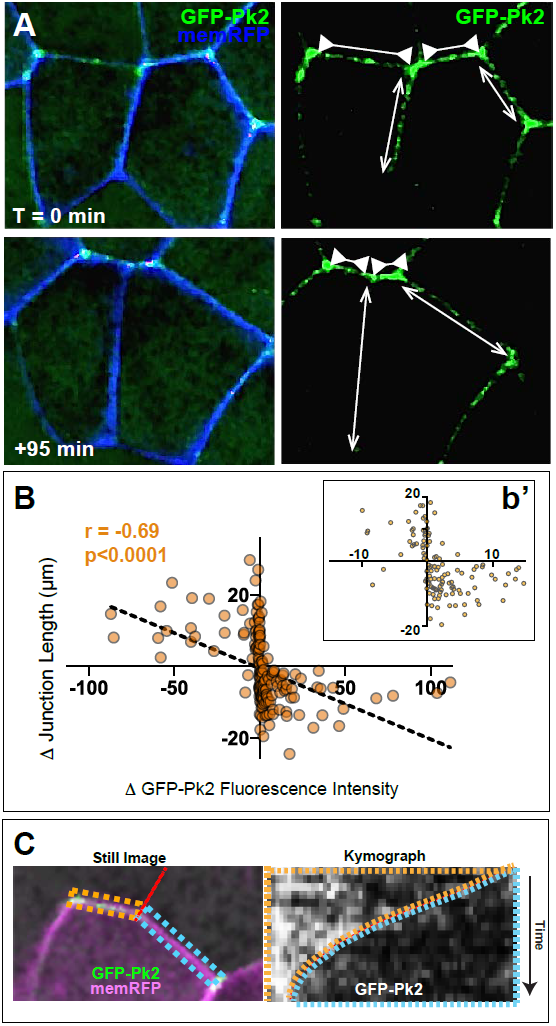
Pk2 is dynamically enriched specifically at shrinking V-junctions. **(A)** Confocal images of neural epithelial cells labeled with membraneRFP and GFP-Pk2 showing the change in length of shrinking junctions (inward facing arrowheads) and growing junctions (outward facing arrows) along with the corresponding change in GFP-Pk2 intensity at two different time points. **(B)** Graph showing the correlation between the change in length and change in mean intensity of GFP-Pk2 for 165 individual junctions. **(b’)** Inset image shows a magnified view of the central core of the plot shown in (B). **(C)** Still image (left) and accompanying kymograph (right) depicting the simultaneous changes in the lengths of a shrinking and a growing cell-cell junction separated by a shared tricellular junction and corresponding changes in GFP-Pk2 fluorescence intensity across these junctions over the course of 4200s. Dashed orange and blue lines indicate the two different junctions and the dashed red line traces the movement of the tricellular junction.

These data suggested that the dynamic enrichment of GFP-Pk2 at anterior cell faces might not simply reflect the junction’s spatial orientation. Additional evidence for this notion arose from the observation that GFP-Pk2 intensities frequently differed radically between adjacent cell-cell junctions, even when these junctions share roughly similar orientations. For example, the still image in Figure 5C shows a single neural epithelial cell expressing GFP-Pk2 (green). This cell borders two unlabeled anterior neighbors (separated by the red line) such that the two junctions indicated by orange and blue dashed boxes both represent “anterior” cell faces. Strikingly, as shown by the kymograph, the junction boxed in orange shrinks over time while the other grows; the GFP-Pk2 signal (white) increases dramatically and specifically in the shrinking junction.

Similar results were obtained for Vangl2, as instantaneous changes in GFP-Vangl2 were also more strongly correlated with concomitant changes in junction length than with spatial orientation (Supp. Fig. 3). For example, Supp. Fig. 3B shows several adjacent cell-cell junctions in two frames of a time-lapse movie, with colored arrows indicating growing or shrinking junctions. In Supp. Fig. 3C, corresponding colors are used to plot GFP-Vangl2 intensity against change in length for each junction. Despite their similar mediolaterally-biased orientations, GFP-Vangl2 intensity dramatically increased at the shrinking junctions (Supp. Fig. 3C, green, orange), while intensity slowly decreases at the adjacent growing junctions (Supp. Fig. 3C, Blue, purple). Membrane-RFP does not display a similar pattern (Supp. Fig. 3C, black dashed lines).

These findings suggest that the strength of asymmetric Pk2 and Vangl2 enrichment at a particular junction is more strongly tied to the dynamic behavior of that junction than it is to the junction’s orientation or to positional information across the tissue. A related observation is that levels of junctional Pk2 and Vangl2 frequently displayed radical changes precisely at tricellular junctions, even if the apposed cell-cell junctions share similar orientation (Fig. 5A; 6A). This result suggests that the enrichment of Pk2 and Vangl2 is governed by the reciprocal interaction of each individual pair of neighboring cells and is intricately linked to the behavior of the shared junction between them.

Finally, we noted that the rate at which GFP-Pk2 and GFP-Vangl2 increased at shrinking junctions often appeared significantly higher than the rate of decrease along growing junctions (e.g. Fig. 6B), suggesting that distinct mechanisms might control the dynamics of PCP protein accumulation at shrinking junctions and loss at growing junctions, as we investigate below. In summary, our data suggest an unexpectedly complex relationship between the spatiotemporal accumulation of PCP proteins and the dynamic behaviors of cell-cell junctions during intercalation.

**Figure 6:**
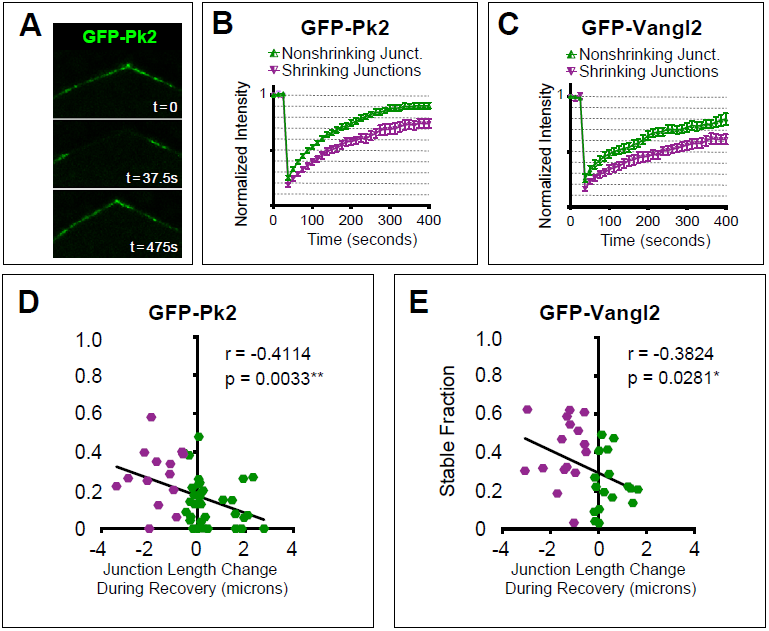
Turnover of Pk2 and Vangl2 correlates with junction behavior. **(A)** Three still images from a time-lapse movie captured before and after photobleaching the anterior apicolateral region of a cell mosaically labeled with GFP-Pk2 in the neural plate. **(B, C)** Graphs showing mean fluorescence recovery after photobleaching at shrinking and nonshrinking junctions for GFP-Pk2 (n=34 nonshrinking, n=15 shrinking) and GFP-Vangl2 (n=17 nonshriking, n=16 shrinking). Shrinking junctions were defined as those that were reduced by 0.5μm or more in length over the course of bleaching and fluorescence intensity recovery. **(D-E)** Graphs plotting the change in junction length during photobleaching and recovery against the calculated nonmobile fraction for the individual junctions analyzed in (B) and (C) with associated linear regression model and correlation analysis statistics included. n=49 (GFP-Prickle2); n=33 (GFP-Vangl2).

**Figure 8:**
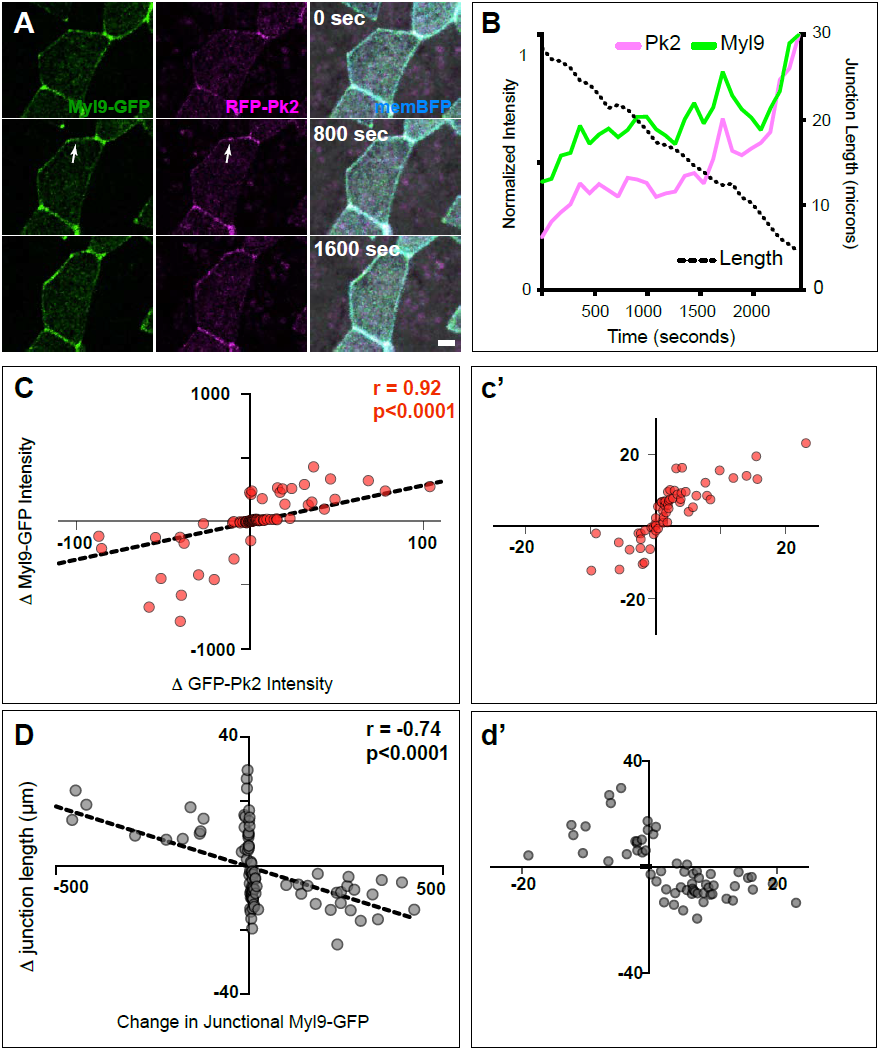
Spatiotemporal coordination of Prickle2 and actomyosin accumulation at shrinking junctions. **(A)** Confocal images of a shrinking AP junction labeled with Myl9-GFP, RFP-Pk2, and membraneBFP over the course of a 1600s time lapse. Note the increased GFP and RFP intensities as the junction becomes shorter. Scale = 10 μm. **(B)** Intensity trace over time for the junction indicted by arrows in panel A and showing pulsed co-accumulation of Pk2 (pink) and Myosin (green) as junction length shrinks (black dashed line). **(C)** Scatter plot correlation of change in RFP-Pk2 intensity plotted against the change in GFP-Myl9 intensity; a magnified view of the core of this plot is shown in Panel **c’**. n=96 junctions, 2 experiments, 3 embryos. **(D)** Scatter plot correlation of change in junction length plotted against the change in GFP-Myl9 intensity; a magnified view of the core of this plot is shown in Panel **d’**. n=46 junctions, 2 experiments, 3 embryos.

### Turnover of Prickle2 and Vangl2 at cell-cell junctions is planar polarized during cell intercalation

Of the PCP protein dynamics we observed, we feel that the enrichment specifically at shrinking V-junctions is the most significant, and we considered two possible interpretations of this observation. First, intensity could reflect density, increasing simply because the amount of protein on a junction remains constant as that junction shrinks. Indeed, generic membrane markers show a tendency to increase in intensity as junctions shrink, but even when normalized against such a membrane label, intensities of Vangl2 and Prickle2 still displayed a significant correlation with junction shrinkage (Supp. Fig. 4). This result prompted us to consider the alternative hypothesis that the observed enrichment could be an active process in which regulated turnover kinetics could be more dynamic at some junctions and less so at others. FRAP studies have demonstrated that polarized PCP protein turnover is planar polarized in other cell types (Butler and Wallingford, 2015; Chien et al., 2015; Shi et al., 2016; Strutt et al., 2011), so we used this method to assess localization dynamics during cell intercalation in the closing neural tube (Fig. 6A).

Both Pk2 and Vangl2 displayed striking differences in turnover kinetics at shrinking versus non-shrinking junctions, with significantly less recovery (i.e. higher stable fraction) for both Vangl2 and Prickle2 at shrinking junctions compared to non-shrinking junctions (Fig. 6B, C). Moreover, when we integrated these FRAP data with time-lapse analysis of cell behaviors, we found that the stable fraction of both proteins correlated significantly with changes in junction length: more rapidly shrinking junctions displayed higher stable fractions of junctional PCP proteins (Fig. 6D, E). The different rates of turnover detected by FRAP are consistent with the results above suggesting that PCP accumulation at shrinking junctions and loss from growing junctions occur at different rates (e.g. Supp. Fig. 3). Thus, the overall accumulation of Prickle2 and Vangl2 at shrinking junctions parallels the increased stability of these proteins at these sites, and together, these data suggest a key role for PCP protein localization in the coordination of cell-cell junction shrinkage.

### Planar polarization of actomyosin during junction shrinking in the neural epithelium is PCP-dependent

We next sought to understand the link between PCP protein localization and the actomyosin machinery known to drive cell-cell junction shrinkage. Time-lapse imaging in *Drosophila* first demonstrated that cell intercalation by junction shrinkage is accompanied by pulsed accumulations of actomyosin at V-junctions (Bertet et al., 2004; Fernandez-Gonzalez et al., 2009; Rauzi et al., 2008). While this process is independent of PCP signaling in *Drosophila*, similar actomyosin pulses have been observed during PCP-dependent junction shrinking in mesenchymal cells of the *Xenopus* gastrula mesoderm (Shindo and Wallingford, 2014). Static analyses of *Xenopus*, chicks and mice also indicate that actomyosin is enriched at mediolaterally oriented junctions (McGreevy et al., 2015; Nishimura et al., 2012; Williams et al., 2014). However, the spatiotemporal relationship between actomyosin dynamics and subcellular behaviors during cell intercalation in the vertebrate neural tube remain poorly defined.

Using a GFP-fusion to the myosin regulatory light chain Myl9 (Shindo and Wallingford, 2014), observed a strong enrichment at V-junctions as compared to T-junctions. This enrichment was apparent at both the population level (Fig. 7A-C, green) and in the significant correlation between of Myl9 intensity and junction angle for individual junctions (Fig. 7D, green, n > 450). Moreover, expression of Xdd1 or Pk2-ΔPΔL significantly disrupted the planar polarization of myosin enrichment (Fig. 7A-D, blue & magenta; n > 240 for each condition). Interestingly, these reagents appear to act via converse mechanisms: Expression of Xdd1 elicited an elevation of myosin levels at T junctions, while expression of Pk2-ΔPΔL elicited a reduction of myosin enrichment at V junctions (Fig. 7C), and a similar trend was observed in the effect of these reagents on GFP-Pk2 intensities (Fig. 1B). Thus, disruption of PCP signaling disrupts both asymmetric PCP protein localization and the coordination of planar polarized actomyosin contraction in the closing neural tube.

**Figure 7:**
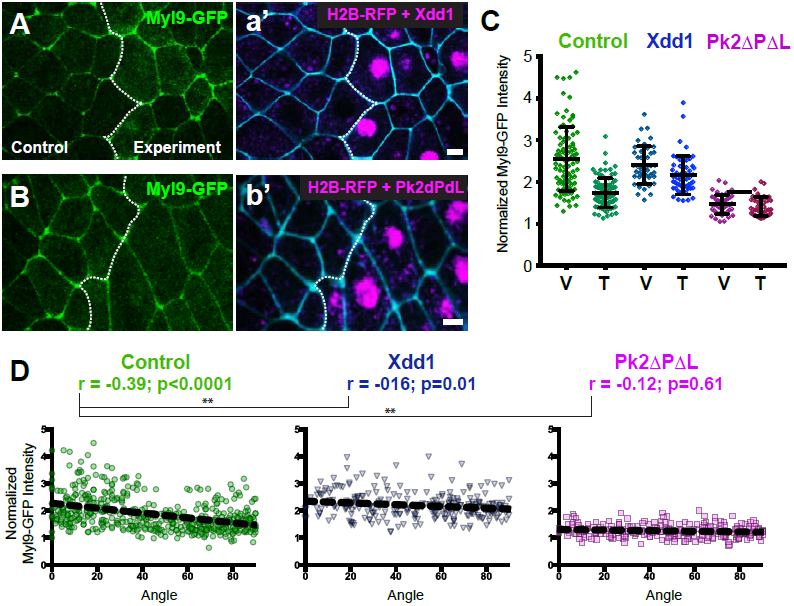
PCP function is required for polarization of actomyosin contractility during junction shrinking. **(A-B)** Confocal images of *Xenopus* neural epithelia labeled evenly with GFP-Pk2 and membraneBFP and mosaically with H2B-RFP serving as a tracer for either Xdd1 (A) or Pk2-ΔPΔL (B) expression. Scale = 10 μm **(C)** Graph plotting Myl9-GFP intensity along V-junctions (0-45° relative to mediolateral axis) and T junctions (46-90° relative to mediolateral axis) normalized as a ratio to the cytoplasmic intensity. Control cells (n=91 V, 91T) and cells expressing Xdd1 (n=44 V, 53 T) and Pk2-ΔPΔL (n=45 V, 45 T). Ctrl V vs. T, p<0.0001****; Pk2-ΔPΔL V vs. T, p=0.2304ns; Xdd1 V vs. T, p=0.0022**; Control V vs. Xdd1 V, p=0.5826ns; Control T vs. Xdd1 T, p <0.0001****; Control T vs. Pk2-ΔPΔL T, p<0.0001**** (Mann-Whitney Test for significance). **(D)** Distributions of normalized Myl9-GFP intensity plotted against the angle of the junction at which intensity was measured in control cells and cells expressing Xdd1 or Pk2-ΔPΔL. Correlation coefficients for Xdd1 and Pk2-ΔPΔL were shown to be significantly different from controls using the Fischer R-to-Z transformation. n=498 junctions (Control), n=263 junctions (Xdd1) and n=245 junctions (Pk2-ΔPΔL) from 4 experiments, 5 embryos (Xdd1); 3 experiments, 7 embryos (Pk2-ΔPΔL).

### PCP proteins and actomyosin dynamics are spatiotemporally coordinated with junction shrinkage

Finally, because of the observed spatiotemporal patterns of PCP protein localization (above), we more closely examined Myl9 localization during time-lapse movies. As expected from results in other systems, Myl9-GFP displayed a pulsatile behavior at shrinking junctions (Fig. 9A, B, green, black lines). Strikingly, Pk2 intensity also displayed pulsatile enrichment, and changes in Pk2 were strongly correlated with similar changes in Myl9 intensity at cell-cell junctions (Fig. 9A, B pink line, C). These correlations are likely to be functionally relevant, because instantaneous changes in myosin enrichment strongly correlated with instantaneous decreases in junction length (Fig. 9D). Similar results were obtained using the actin biosensor Utr-RFP (Burkel et al., 2007), and the changes in myosin intensity were also strongly correlated to changes in actin on individual junctions (Supp. Fig. 5). Together, these data demonstrate shared dynamic patterns of core PCP protein and actomyosin localization in time and space during cell intercalation and suggest that these systems work in close concert to drive junction shrinking in the *Xenopus* neural epithelium.

## Discussion

Here, we have used image tiling time-lapse microscopy in *Xenopus* embryos to generate high magnification movies of the closing vertebrate neural tube. These movies allowed us to quantify protein localization and dynamics as they relate to the cell behaviors associated with convergent extension. We focus here on junction shrinkage, which together with mediolateral protrusions is an essential sub-cellular behavior contributing to cell intercalation. We find that the core PCP proteins Prickle2 and Vangl2 display a consistent pattern of localization and turnover in space and time during cell intercalation.

The spatial asymmetry of core PCP proteins is fundamental to their function. In a wide array of cell types, planar polarization is defined by Dvl and Frizzled enrichment to one region of the cell and Vangl and Prickle in a reciprocal pattern. Feedback across cell membranes is thought to reinforce initially weak asymmetry leading to the robust asymmetry at later stages, as observed first in *Drosophila* (Butler and Wallingford, 2017; Strutt and Strutt, 2009).

Interestingly, while cell intercalation was the first setting in which vertebrate PCP was defined, the first reports of asymmetric protein localization in vertebrates came instead from studies of cochlear hair cells and later from ciliated cells of the node and airway (Rida and Chen, 2009; Wallingford, 2010). Indeed, even now we know little about the spatial patterns and almost nothing about the temporal patterns of PCP protein localization in the context of convergent extension. Given the key role for PCP-dependent convergent extension in human neural tube defects (Wallingford et al., 2013), we feel that the data here -though largely descriptive- are nonetheless highly significant.

Currently, a consensus has emerged that PCP proteins spatially delineate an anterior-to-posterior axis in cells undergoing cell intercalation. This consensus has support from studies of various PCP proteins in diverse tissues, first in zebrafish and later in other animals (Ciruna et al., 2006; McGreevy et al., 2015; Nishimura et al., 2012; Ossipova et al., 2015; Roszko et al., 2015; Yin et al., 2008). This model is consistent with the similar pattern of anteroposterior localization for PCP proteins in the orientation of directional ciliary beating in the embryonic node and spinal cord (Antic et al., 2010; Borovina et al., 2010; Hashimoto et al., 2010). Moreover, this axis of polarization is consistent with embryological data suggesting that anteroposterior patterning is crucial to convergent extension in *Xenopus* (Ninomiya et al., 2004). However, it should be noted that several studies suggest additional regions of localization in the mediolateral ends of cells during cell intercalation (e.g. (Jiang et al., 2005; Kinoshita et al., 2003; Panousopoulou et al., 2013)).

Our data here are consistent with the idea of anteroposterior localization of PCP proteins plays a critical role in cell intercalation, though they do not exclude additional roles in mediolateral protrusions, and disruption of PCP does disrupt the polarity and stability of mediolateral protrusions (Wallingford et al., 2000). Significantly, recent work suggests that both mediolateral protrusions and junction shrinking act together during convergent extension (Sun et al., 2017; Williams et al., 2014), so it is clear that additional studies will be required.

More important than the spatial localization of PCP proteins during cell intercalation is the temporal aspect reported here, as previous studies have provided only static snapshots of what is a highly dynamic process. Our time-lapse studies not only reveal that Prickle and Vangl are dynamically enriched at shrinking junctions, but also suggest that the shrinking/growing status of a junction is a better indicator of PCP protein enrichment than is its orientation in the mediolateral/anteroposterior axis. In addition, our FRAP data reveal that turnover kinetics also differ with the dynamic behavior of the junctions, with higher stable fractions associated with shrinking junctions. Thus, our study reveals a new complexity to the pattern of PCP protein localization that likely reflects the dynamic nature of the cells involved, as they are constantly exchanging neighbors as they intercalate.

Another interesting result in this study is the tight spatiotemporal relationship between PCP proteins and myosin at shrinking junctions. PCP proteins were first found to drive the phosphorylation of Myosin in *Drosophila* (Winter et al., 2001), and more recently the same has been shown for cell intercalation in *Xenopus* and mice (Lienkamp et al., 2012; McGreevy et al., 2015; Nishimura et al., 2012; Shindo and Wallingford, 2014; Williams et al., 2014), as well as in the cochlea of mice (Lee et al., 2012). Interestingly, two recent studies in *Xenopus* and ascidians suggest that myosin also acts upstream of PCP protein localization (Newman-Smith et al., 2015; Ossipova et al., 2015) and may involve a reciprocal relationship in their polarized functions, and this notion is supported by our data here.

One limitation to our study is the focus here on Prickle and Vangl. Attempts to perform similar experiments with Dvl and Frizzled proved too challenging, but of the PCP proteins it is Dvl that is most directly linked to Myosin activation, as Dvl binds to and activates RhoA via Daam1 (Habas et al., 2001), which can act via Rho kinase to phosphorylate myosin. However, less is known about how Myosin dynamics are controlled on the opposing, Prickle/Vangl-enriched cell face, so this remains an important question to answer. One attractive hypothesis involves Dvl/RhoA-mediated myosin accumulation on one cell face resulting in mechanically-induced myosin accumulation on the opposing cell face. Indeed, mechanical positive feedback has been shown to contribute to the oscillations of actomyosin at shrinking junctions during PCP-independent cell intercalation in *Drosophila* (e.g. (Fernandez-Gonzalez et al., 2009)).

Finally, our data provide an interesting complement to previous studies linking turnover at cell junctions to planar polarized PCP protein localization. In *Drosophila*, the enriched regions of PCP protein localization display higher stable fractions than do non-enriched regions, and the loss of asymmetric protein localization after disrupting PCP signaling is accompanied by a loss of this asymmetric turnover (Strutt et al., 2011). Similar results have been obtained in vertebrate multiciliated cells on the *Xenopus* epidermis and in the mouse oviduct {Butler, 2015 #6091;Chien, 2015 #6498}. However, cell-cell neighbor exchange is minimal in these contexts, in contrast to cell intercalation, where such neighbor exchanges are constant. We therefore find it interesting that we observed a similar trend during cell intercalation, where enriched regions of PCP protein localization also display higher stable fractions of PCP protein even as these junctions shrink dramatically. Interestingly, when we calculated the mean stable fraction for PCP proteins (i.e. averaging all junctions), this value was substantially lower than what we previously observed using similar methods in *Xenopus* multiciliated cells (Butler and Wallingford, 2015); this difference may represent an adaptation to the dynamic nature of the junctions involved. Together with the previous studies, our data demonstrate that planar polarized junction turnover kinetics are a general feature of PCP protein localization, spanning a broad spectrum of organisms and cell types. The data here thus provide a quantitative, dynamic view of PCP protein localization as it relates to a subcellular behavior that drives cell intercalation and lays a foundation for further study of PCP protein function during neural tube closure.

## Materials and Methods

### *Xenopus* manipulations

Eggs were and externally fertilized according to standard protocols (Sive et al., 2000). The jelly coat was removed from embryos at the 2-cell stage by bathing in a solution of 2% cysteine (pH 7.9). The embryos were then washed in 1/3x Marc’s Modified Ringer’s (MMR) solution and microinjected in a solution of 1/3x MMR with 2% Ficoll using an Oxford manipulator. The mRNAs coding for fluorescent protein fusions were synthesized using mMessage mMachine kits (Ambion) and injected into one of eight dorsal blastomeres for even labeling of the neural plate at the following concentrations: 50pg membraneGFP, 60pg of membraneRFP, 80pg of membraneBFP, 200pg for GFP- or RFP-Pk2, 60pg for GFP-Vangl2, 50pg for H2B-RFP, and 30pg for Myl9-GFP. For mosaically labeled tissues, mRNAs were injected at the 16- or 32-cell stages with approximately 70% of the totals amounts listed above. Dominant-negative Pk2 (Pk2-ΔPETΔLIM) was injected at 700-800pg for overexpression, at the eight-cell stage, as was the dominant-negative Dvl (Xdd1), and both were similarly reduced by 70% for later stage injections. For Pk2 morpholino treatments, 20-25ng was injected into one cell at the eight-cell stage, and 400pg of GFP-Pk2 were used to rescue the morpholino phenotypes. Developmental stages were determined according to (Nieuwkoop and Faber, 1994).

### Live Imaging and Image Quantification

Confocal imaging was done using live embryos submerged in 1/3x MMR in AttoFluor Cell Chambers (Life Technologies A7816) and between coverglasses, using silicon grease as an adhesive spacer, and carried out with a Zeiss LSM700 confocal microscope. Images were processed with the Fiji distribution of ImageJ, Imaris (Bitplane) and Photoshop (Adobe) software suites, and figures were assembled in Illustrator (Adobe). For junction length and protein enrichment measures, freehand lines (3-6 pixels wide, depending on image scale/zoom) were drawn over anterior cell membranes, while cytoplasmic measures were taken with the freehand shapes tools. Statistical analyses were carried out using Prism (Graphpad) software with Mann Whitney tests for significance and Spearman non-parametric correlations. Extreme outliers with fluorescent intensities more than three standard deviations away from the mean were removed from the analysis; in all datasets such outliers were very rare (<6). These outliers likely represent non-specific contraction waves sometimes observed in the *Xenopus* neural plate. For FRAP analysis, time-lapse movies were acquired after photobleaching discrete domains of core PCP GFP fusions localization. Intensity measurements were taken in Fiji, with recordings for each time point taken individually from each frame captured at bleached regions and normalized as detailed in (Goldman RD et al., 2005). Statistical analysis was performed in Prism (Graphpad) software with exponential decay functions. Angles of junctions shrinking to form and emerging from rosettes were measured manually in Fiji, and rose diagrams were plotted with Oriana software (Kovach Computing Services). Stereo time-lapse imaging was performed using a Zeiss AXIO Zoom.V16 Stereomicroscope and associated Zen software. Movies were exported from Fiji and processed in Adobe Photoshop, and cell tracking was performed with the Fiji manual tracking plug-in.

## Supplemental Figure Legends

**Supp. Fig. S1:**
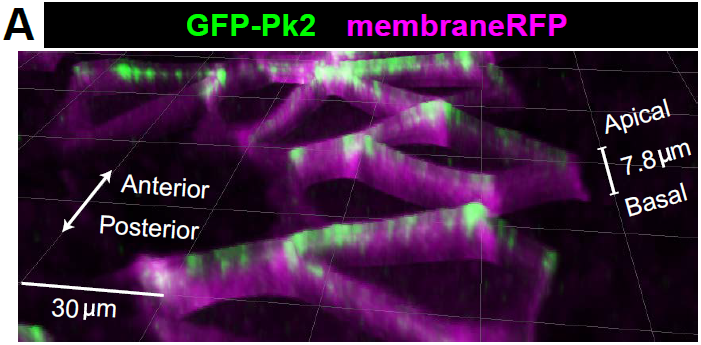
GFP-Pk2 localizes to anterior apicolateral regions of cells in the *Xenopus* neural plate. Pseudo 3D render of neural epithelium mosaically labeled with GFP-Pk2 and membraneRFP showing Pk2 restricted to the apicolateral cellular regions.

**Supp. Fig. S2:**
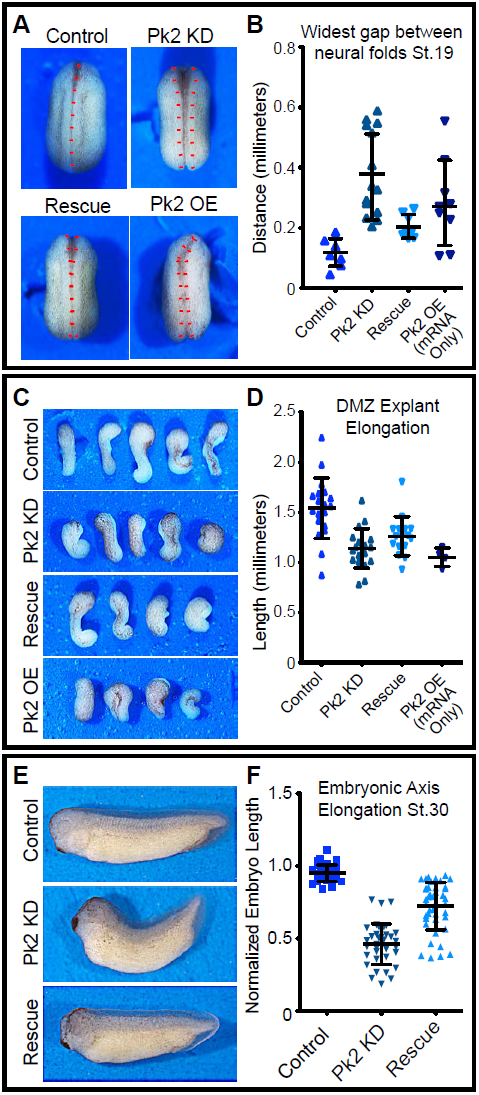
Pk2 knockdown results in embryonic convergent extension phenotypes. **(A)** Dorsal view Stereoscope images of a representative Stage 20 control embryo, an embryo that has been injected dorsally with 25ng Pk2 morpholino, an embryo that received morpholino plus a 400pg GFP-Pk2 mRNA rescue dose, and an embryo that received only the 400pg Pk2 mRNA. Anterior is up **(B)** Graph of the distance between the neural folds at stage 20. n=8 (Control), 14 (Pk2 KD), 8 (Rescue), 9 (RNA/OE). Control vs. KD, p<0.0001; vs. Rescue, p=0.0019; vs. OE, p=0.0079. (Mann-Whitney Test for significance) **(C)** Images of Dorsal marginal zone (DMZ) explants dissected from stage 11.5 embryos of the treated under the same conditions as in (**A)**.**(D)** Graph of DMZ explant length at a time when embryos that had not been dissected had reached Stage 20. n=19 (Control), 19 (Pk2 KD), 16 (Rescue), 4 (RNA/OE). Control vs. KD, p<0.0001; vs. Rescue, p=0.0010; vs. OE, p=0.0041. (Mann-Whitney Test for significance) **(E)** Representative images of Stage 30 embryos that received 30ng Pk2 morpholino, morpholino plus 400pg rescue GFP-Pk2 mRNA, or controls that received neither. Anterior is left and dorsal is up. **(F)** Graph of the length of the dorsal embryonic length of embryos represented by those shown in **(E)**. n=35 (Control), n=37 (Pk2 KD), n=44 (Rescue). Control vs. KD, p<0.0001; vs. Rescue, p<0.0001. (Mann-Whitney Test for significance)

**Supp. Figure S3:**
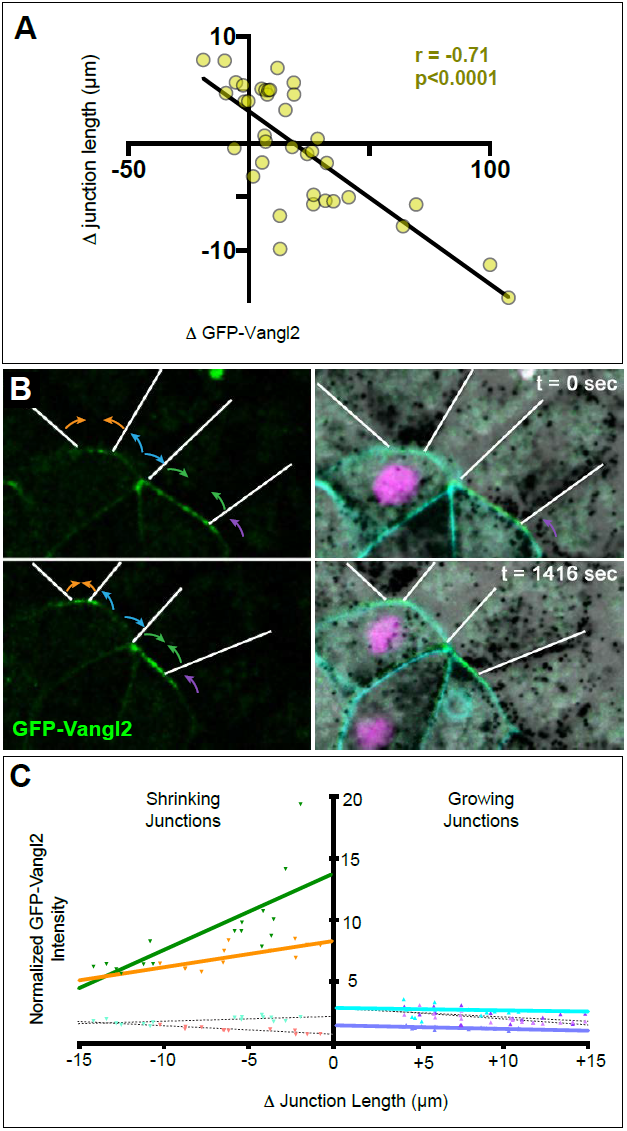
Distinct Vangl2 dynamics at adjacent growing and shrinking junctions. **(A)** Confocal stills of a neural epithelium mosaically labeled with GFP-Vangl2 (left) and co-labeled with H2B-RFP and membraneBFP (right). White lines indicate junctions of unlabeled neighboring cells. Colored arrows indicate shrinking (orange, green) or growing (blue, purple) behaviors of adjoined cell-cell junctions. Note dramatic changes of Vangl2 intensity at tricellular junctions. **(B)** Graph plotting the changes in GFP-Vangl2 (solid lines) intensity at the correspondingly colored junctions shown in (A). Note robust accumulation at shrinking junctions and modest loss at growing junctions despite roughly similar orientations. MembraneBFP levels (dashed lines) do not change significantly. Lines in (B) represent fits of a linear regression model.

**Supp. Figure S4:**
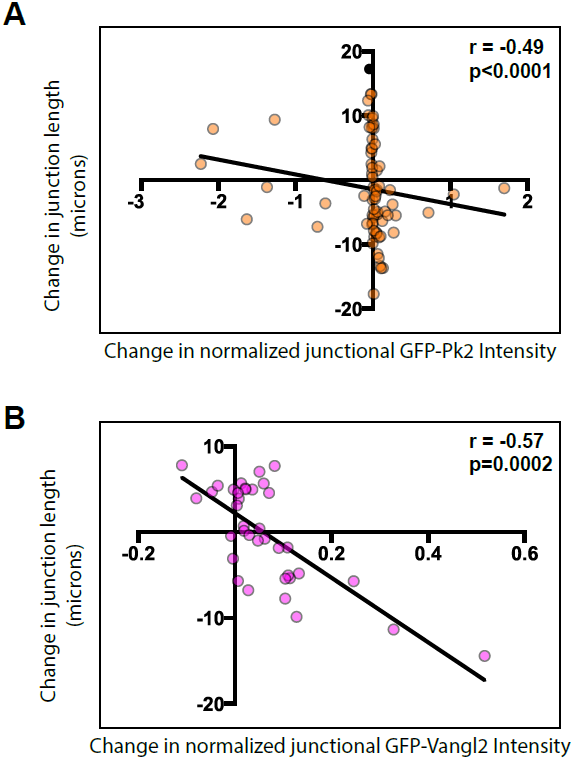
**(A).** Plot of normalized GFP-Pk2 intensities versus change in junction length (r = -0.49; p<0.001; n=71). Normalization was performed by subtracting background and expressing the intensity of GFP-Pk2 levels as the ratio to the intensity of co-expressed membrane-BFP. (**B)** Plot of normalized GFP-Vangl2 intensities versus change in junction length (r = -0.57; p=0.002; n=37). Normalization was performed as per GFP-Pk2, above.

**Supp. Figure S5:**
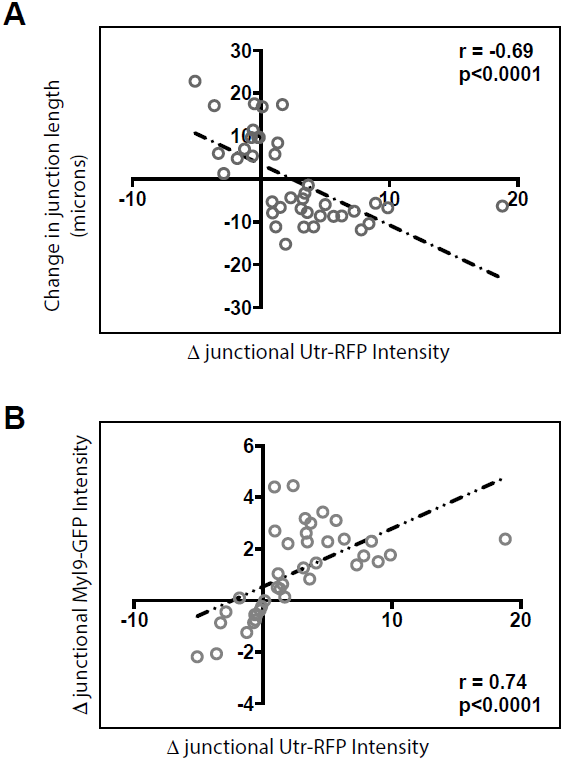
**(A)** Plot showing the correlation between changes in junction length and Utrophin-RFP (actin biosensor) intensities. **(B)** Plot showing the correlation between changes in Myl9-GFP and Utrophin-RFP (actin biosensor) intensities. n=39 junctions for each plot.

